# Disruption of Hepatocyte *Jak2* leads to Spontaneous NASH in Aged Mice and Uncouples Metabolic Liver Disease from Insulin Resistance

**DOI:** 10.1101/079236

**Authors:** Camella G. Wilson, Aras N. Mattis, Jennifer L. Tran, Kevin Corbit, Ethan J. Weiss

## Abstract

Growth Hormone (GH) is a master regulator of metabolic homeostasis and longevity. Whole body GH insensitivity (GHI) augments insulin sensitivity, age-related disease resistance, adiposity, and occurrence of NAFLD. Conversely, acromegalic patients are prone to diabetes and increased mortality due to constitutive high levels of circulating GH. However, which tissues control the various metabolic aspects of GH physiology are unknown. Therefore, we determined the role of GH in age-related metabolic dysfunction by inducing hepatocyte- (JAK2L) or adipocyte-specific (JAK2A) GHI individually or combinatorially (JAK2LA) via deletion of *Jak2*, an obligate transducer of GH signaling. Aged JAK2L mice were insulin resistant but lean and had significant NASH, hepatic inflammation, and fibrosis. In contrast, JAK2A animals had increased adiposity and were completely resistant to age-associated hepatic steatosis, NASH, and insulin resistance. Interestingly, while JAK2LA mice retained enhanced whole-body insulin sensitivity, they still developed NASH to an almost identical degree as JAK2L mice but with a substantial reduction in the degree of microvesicular steatosis. Collectively, loss of adipocyte *Jak2* conferred whole body insulin sensitivity even in the face of obesity and NASH. Deletion of hepatocyte *Jak2* promoted NASH in aged mice without any dietary or drugs perturbations. The effect appears to be liver autonomous and cannot be overcome by the insulin sensitizing effect of adipocyte *Jak2* deletion. Here, we describe the first model of spontaneous NASH that is coupled to augmented insulin sensitivity. Further, there was an inverse correlation between insulin sensitivity and the degree of microvesicular steatosis. Therefore, GH signaling independently mediates insulin/glucose and lipid homeostasis and directly regulates the development of NASH in aged mice.

**Financial Support:** This study was supported by National Institutes of Health (NIH) Grants 1R01DK091276 (to E.J.W.). We also acknowledge the support of the University of California, San Francisco (UCSF) Cardiovascular Research Institute, the UCSF Diabetes Center (P30 DK063720), the UCSF Liver Center (P30 DK026743, and the James Peter Read Foundation.

**Abbreviations:**

NASH: non-alcoholic steato-hepatitis
NAFLD: non-alcoholic fatty liver disease
GH: growth hormone
JAK2: Janus kinase 2
CON: CON mice
JAK2L: hepatocyte-specific deletion of JAK2
JAK2A: adipocyte-specific deletion of JAK2
JAK2LA: hepatocyte and adipocyte JAK2 knockout
TG: triglyceride
AST: aspartate aminotransferase
ALT: alanine transaminase
Stat5: signal transducer and activator of transcription 5
qRT-PCR: quantitative reverse-transcription polymerase chain reaction
Mcp1: monocyte chemoattractant protein-1
Cd11b: cluster of differentiation molecule 11b
F4/80: EGF-like module-containing mucin-like hormone receptor-like 1
FcgR1: high affinity immunoglobulin gamma Fc receptor I
L-Fabp: liver fatty acid binding protein
PPARγ: peroxisome proliferator-activated receptor gamma
FATP: fatty acid transport protein
CD36/FAT: Fatty Acid Translocase
ITT: insulin tolerance test.
Lpl: lipoprotein lipase
IL-: interleukin-
FcgR1: Fc receptor IgG
Tnfα: tumor necrosis factor alpha
Tgfβ1: transforming growth factor beta 1
αSMA, alpha 2: smooth muscle actin
IGF-1: insulin-like growth factor 1.

## INTRODUCTION

Non-alcoholic fatty liver disease (NAFLD) is highly prevalent, and its growth parallels rising epidemics of obesity and type 2 diabetes mellitus (T2DM), and it is considered the hepatic manifestation of metabolic syndrome (1). Within the spectrum of NAFLD, simple steatosis (fatty liver) is thought to be relatively benign. However, in 10% to 20% of patients, steatosis progresses to nonalcoholic steatohepatitis (NASH), where excess liver fat is coupled with inflammation and liver cell damage that can progress to cirrhosis and liver failure (2, 3). Risk factors for NASH progression include age, obesity, insulin resistance, growth hormone deficiency and viral hepatitis (4, 5).

Liver lipid homeostasis results from a proper balance of lipogenesis, uptake, secretion, oxidation and lipolysis, and derangements within those can contribute to the initiation and progression of NAFLD. Indeed, patients with NAFLD have augmented fatty acid synthesis (7). Increased hepatic fatty acid uptake, which occurs mainly through the SLC27A transport proteins and the scavenger receptor CD36, can also lead to NAFLD. The mechanism of this dissociation between fatty liver and insulin resistance is unknown, but it has been reported that free fatty acids (FFA) and not hepatic triglyceride (8) are associated with NAFLD (9). Interestingly, the rate of adipose tissue lipolysis, which produces FFA, is increased in NAFLD and NASH (10, 11).

Recently, adipose tissue lipolysis has been identified as a major regulator of hepatic insulin sensitivity (12, 13). This work stems from the surprising finding that insulin is able to suppress hepatic glucose output in the absence of liver insulin signaling (14, 15). Therefore, adipose tissue mediates some of the metabolic consequences in liver, including NAFLD and NASH (15, 16). Indeed, hepatic insulin resistance strongly correlates with and has a possible causative role in the development of NAFLD (17). However, there are examples of NAFLD and NASH uncoupled with insulin resistance. Variants of *PNPLA3* and *TM6SF2* associated with fatty liver maintain adipocyte (i.e. inhibition of lipolysis) and hepatocyte (i.e. inhibition of glucose production) insulin sensitivity (18). Moreover, knockdown on *CGI-58*, the Chanarin-Dorfman syndrome gene, promotes advanced NAFLD yet protects from high fat diet associated hepatic insulin resistance (19). Therefore, the cause-and-effect relationship of insulin resistance and NAFLD is not entirely clear.

GH is known to regulate adipose tissue lipolysis and hepatic lipid homeostasis (20, 21) and is known to be involved in NAFLD (22). Several groups have reported mouse models demonstrating the development of NAFLD as the result of impaired hepatic GH signaling (23, 24). Disruption of GH signaling via deletion of *Stat5* or *Ghr* results in hepatic steatosis, fibrosis and liver cancer (24, 25). Further, GH deficiency is associated with increased risk for NAFLD in adult humans (26, 27). In the JAK2L model, where GH signal transduction is selectively disrupted in hepatocytes, a redistribution of fat from adipose tissue to the liver is appreciated resulting in up to a 20-fold increase in liver TG and cholesterol in 8-10 week old mice (21, 28). In a 20-week old cohort of these mice, early signs of hepatic inflammation were present, but without frank NASH.

There are few examples of spontaneous NASH developing in rodents in the absence of special diets or other chemical interventions. Studies using the methionine-choline deficient diet and the high fat and/or carbohydrate diet have highlighted the roles of excess hepatic fatty acid uptake, oxidative stress and liver cell injury in disease development (29-31). Genetic models of impaired leptin signaling and Srebp deletion have demonstrated how obesity, insulin resistance and immune response contribute to the disease (32, 33). Though these models do not exactly recapitulate the human presentation of NASH, several of the key features of NASH histopathology are shared (34, 35)

We sought to determine whether aged JAK2L mice would eventually develop overt NASH. To do so, we studied a 40-week cohort of the original mixed background JAK2L mice and we found that hepatic steatosis indeed progressed to NASH in mice with hepatic *Jak2* deletion. Previously, we reported that deleting *Jak2* from adipocytes in JAK2L mice rescues hepatic steatosis in what we termed, JAK2LA mice (36). This led us to consider whether JAK2LA mice might be protected from NASH as compared to JAK2L mice. Therefore, we studied a second independent cohort of mice in an inbred C57BL/6 background at 70-weeks and found that deletion of *Jak2* from adipocytes protects aged JAK2A mice from aging-associated hepatic steatosis, NASH, and insulin resistance. In contrast, JAK2LA mice had only a mild protection from simple steatosis and were not protected at all from developing NASH. Interestingly, despite development of NASH, JAK2LA mice retained whole-body insulin sensitivity. Therefore, the development of NASH appears to be liver autonomous and regulated by loss of hepatic *Jak2* signaling. And critically, for the first time, we have demonstrated a mouse model of spontaneous, diet-independent NASH that is dissociated from the changes in body composition and insulin sensitivity.

## MATERIALS AND METHODS

### Animals

Male Control (CON) and JAK2L mice on C57BL/6,129/Sv background were generated as previously described (21) and maintained for 40 weeks. CON, JAK2L, JAK2A and JAK2LA mice were bred as previously described (36) and maintained for 65-70 weeks. All mice were housed in a constant 12-h light/dark cycle and fed chow diet with 22% caloric intake from fat (Pico Lab Diet #5058 St Louis, Missouri) and water *ad libitum*. All animal studies were performed in compliance with the approved protocols of the University of California San Francisco Institutional Animal Care and Use Committee.

### DXA and Tissue collection

Lean and fat mass were determined using dual-energy x-ray absorptiometry (DXA) on isoflurane-anesthetized mice. Mice were scanned on a Lunar PIXImus2 densitometer (GE Medical Systems, Fairfield, Connecticut). For final tissue collection, samples were collected from *ad-libitum* fed mice starting at 10AM using standard sterile techniques. Tissue was harvested and immediately snap frozen in liquid nitrogen or suspended in 4% PFA for histological analysis.

### Measurement of liver and serum metabolites

For liver triglyceride and cholesterol measurements, tissue was first homogenized in TG buffer (250 mM sucrose, 50 mM Tris, pH 7.4) prior to colorimetric detection using Infinity triglyceride reagent (Thermo Scientific, Middletown VA) or Cholesterol E kit (Wako Chemicals, Richmond VA) respectively. Non-esterified fatty acids (NEFA) esters were measured using commercially available kits (Wako Chemicals). Glucose levels were measured with Contour® glucometer (Bayer, Whippany NJ) via tail vein blood. Blood was collected from the retro-orbital sinus for final blood collection or via the tail vein for fasting and re-fed samples and spun at 15K x g for plasma collection or allowed to clot prior to centrifugation at 10K x g for 10 minutes at 4 °C for serum collection. Plasma and serum were stored at –80 °C until used for analyses. For serum insulin measurements, mice were fasted overnight (16 hours) or fed *ad libitum* prior to blood collection at 10AM. Insulin levels were measured by ELISA (ALPCO Diagnostics, Salem NH) according to the manufacturer’s instructions. Serum triglycerides, cholesterol, HDL cholesterol, LDL cholesterol and liver and kidney function markers (aspartate aminotransferase (AST), alanine aminotransferase (ALT), alkaline phosphatase, bilirubin, creatinine, blood urea nitrogen, albumin and total protein) were analyzed by the UC Davis Comparative Pathology Lab, Davis CA. Serum levels of inflammatory cytokines were measured using the Bio-Plex Pro™ Mouse Cytokine, Chemokine, and Growth Factor Assays (Bio-Rad, Hercules CA) and quantified using a MAGPIX machine (Luminex, Austin TX).

### RNA Isolation and real-time qRT-PCR

We homogenized 30 mg of frozen liver in TRIzol reagent (Invitrogen, Carlsbad CA) and total liver RNA extracted with the RNeasy Mini Kit (QIAGEN Inc.) as previously described (37). TaqMan real-time qRT-PCR was performed with SensiFAST Probe One-Step kit (Bioline, Tauton MA) using 40-50 ηg total RNA for each sample in duplicate. Assays were run on the ABI 7900HT system (Applied Biosystems, Foster City CA). Expression levels were normalized to either ribosomal protein S9 or 18S rRNA and relative levels compared to control samples using the 2-∆∆Ct method (8).

### Liver Histology and Pathology

Formalin-fixed liver tissue was dehydrated in 70% ethanol prior to embedding in paraffin. Sectioning and haematoxylin and eosin and Masson’s Trichrome staining were performed by AML Laboratories Inc (Baltimore, MD). Slides were scored blindly for NASH criteria based on Brunt staging and scoring criteria (38). Images were taken on an Olympus BX51 microscope and edited in Photoshop CC (Adobe, San Jose CA.

### Metabolic tests and fasting experiments

Fasting glucose and insulin levels were collected from overnight (16hrs) fasted mice. For insulin tolerance tests (ITT), mice were fasted for 4 hours then given an i.p. bolus of 1.0U insulin (Novolin® Novo Nordisk, Bagsvaerd Denmark) per kilogram of body weight. Glucose levels at indicated time points were measured from tail vein blood as described above.

### Statistical analysis

Data are presented as means ± SEM. Statistical significance was determined by unpaired two-tailed Student’s *t* test or ANOVA using Prism (Graph Pad, La Jolla, CA). A P value of <0.05 was considered statistically significant.

## RESULTS

### NAFLD in JAK2L mice progresses to NASH with aging

Mice with liver-specific deletion of *Jak2* (JAK2L) develop severe hepatic steatosis starting at 3 weeks of age due to impaired growth hormone (GH) signaling in the liver (21). In a 20-week old cohort of JAK2L mice on mixed C57BL/6,129/Sv background, we observed inflammatory foci and collagen deposition but not frank NASH (Sos et al Supplemental Figure S1). To address if steatosis in these mice would progress to NASH, we maintained this original cohort of mixed background mice on a standard chow diet for 40 weeks then collected tissues for histological and biochemical analyses.

Haematoxylin and eosin (H&E) staining of liver sections showed the presence of hepatic steatosis in both control and JAK2L mice (Figure 1A) but JAK2L mice had 90% of hepatocytes with lipid droplets as compared to 60% of control hepatocytes (Figure 1B right). In both groups, steatosis was localized to zones 3 and 2 but the type of steatosis was markedly different between the two groups with the JAK2L cohort having more macrovesicular and less microvesicular steatosis than control animals (Figure 1B left). Lipid distribution in NASH patients is predominantly macrovesicular and often starts in zone 3 (2).

**Figure 1.**
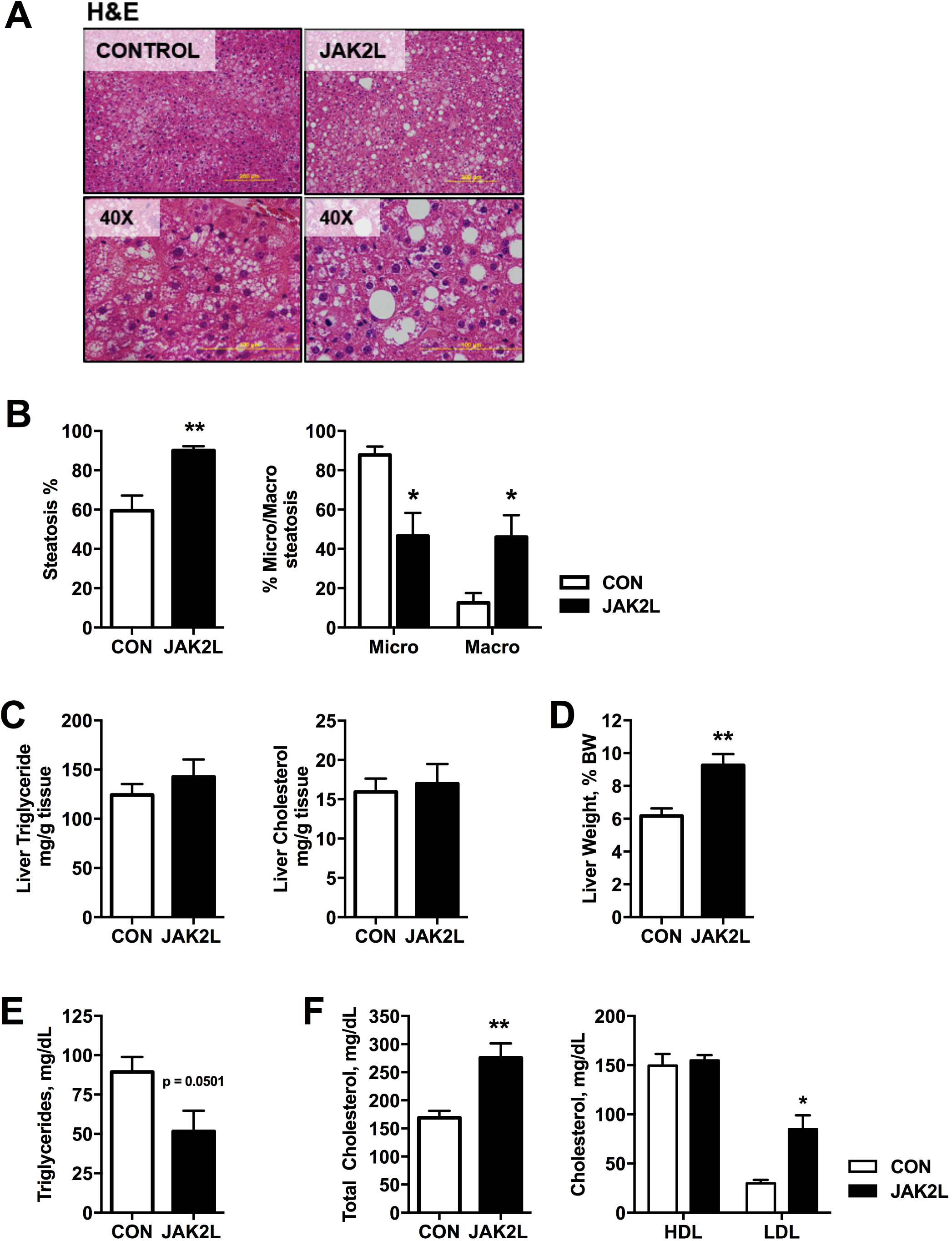
Age induced hepatic steatosis is more severe in JAK2L mice. Control and JAK2L mice on C57BL/6,129/Sv background were maintained on chow diet for 40 weeks prior to assessing liver lipid content. (A) Hematoxylin and eosin (H&E) stain of Control and JAK2L liver at 10X and 40X magnification. (B) Total steatosis and micro-vesicular and macro-vesicular steatosis percentages were assessed from liver H&E sections. (C) Liver triglyceride and cholesterol levels in ad libitum fed mice. (D) Liver weight expressed as a percent of total body weight. (E) Serum triglycerides and (F) cholesterol levels from blood samples collected on day of tissue collection. Data represent mean ± SEM, n=8 for Control and n=5 for JAK2L. * P < 0.05, ** P < 0.01

Surprisingly, liver TG and cholesterol levels were not significantly different between the groups (Figures 1C). Previous reports showed significantly less liver TG in 8-week old control mice compared to this older cohort (21, 28) suggesting that some degree of excess hepatic lipid accumulation occurs spontaneously with aging in this background and on this diet. Liver weights were similar between the cohorts (Table 1) but when normalized to body weight, liver weight percent was 9.25 ± 0.69 % in JAK2L mice compared to 6.17 ± 0.46 % in control animals (Figure 1D). This is not surprising given the lower body mass of the JAK2L cohort.

**Table 1:** Percent of Control and JAK2L mice with histological markers of liver inflammation.

| **Marker** | **Control**<br>**N=8** | **JAK2L**<br>**N=5** |
| --- | --- | --- |
| **Lobular Inflammation** | **%** | **%** |
| No foci | 50.0 | 0 |
| <2 foci per 200X field | 37.5 | 20.0 |
| 2-4 foci per 200X field | 12.5 | 40.0 |
| >4 foci per 200X field | 0 | 40.0 |
| **Microgranulomas** |  |  |
| Present | 25.0 | 80.0 |
| **Large Lipogranulomas** |  |  |
| Present | 0 | 0 |
| **Portal Inflammation** |  |  |
| None to minimal | 87.5 | 40.0 |
| Greater than minimal | 12.5 | 60.0 |
| **Liver Cell Injury** |  |  |
| Few Balloon cells | 12.5 | 20.0 |
| Acidophil bodies | 12.5 | 40.0 |
| **Pigmented Macrophages** | **25.0** | **80.0** |

**Table 2:** Histological Markers of Inflammation in Control, JAK2L, JAK2A and JAK2LA mice.

| **Marker** | **Control**<br>**N=5** | **JAK2L**<br>**N=5** | **JAK2A**<br>**N=5** | **JAK2LA**<br>**N=5** |
| --- | --- | --- | --- | --- |
| **Lobular Inflammation** | % | % | % | % |
| No foci | 0 | 0 | 40 | 0 |
| <2 foci per 200X field | 80 | 80 | 60 | 40 |
| 2-4 foci per 200X field | 20 | 0 | 0 | 60 |
| >4 foci per 200X field | 0 | 20 | 0 | 0 |
| **Microgranulomas** |  |  |  |  |
| Present | 60 | 100 | 20 | 100 |
| **Large Lipogranulomas** |  |  |  |  |
| Present | 40 | 20 | 0 | 40 |
| **Portal Inflammation** |  |  |  |  |
| None to minimal | 80 | 80 | 80 | 60 |
| Greater than minimal | 20 | 20 | 20 | 40 |
| **Liver Cell Injury** |  |  |  |  |
| Few balloon cells | 60 | 80 | 0 | 60 |
| Many balloon cells | 0 | 0 | 0 | 20 |
| **Pigmented Macrophages** | 0 | 20 | 0 | 20 |

Circulating TG levels were lower in JAK2L mice as compared to controls (Figure 1E). On the other hand, total and LDL cholesterol were significantly elevated in the JAK2L cohort (Figure 1F). These data demonstrate that while hepatic steatosis develops spontaneously with age in these mice, JAK2L mice have more pervasive steatosis that is associated with elevated LDL cholesterol levels.

### Hepatic inflammation is increased in aged JAK2L livers

In addition to steatosis, critical features of NASH are hepatic inflammation and liver fibrosis (3). Therefore we assessed the inflammatory state of the liver in the aged mice. Alanine aminotransferase (ALT) and aspartate aminotransferase (AST) are serological markers of liver damage and inflammation and were significantly elevated in JAK2L as compared to control mice (Figure 2A). Alkaline phosphatase, blood urea nitrogen and bilirubin levels were also elevated in JAK2L mice (Supplemental Table 2).

**Figure 2.**
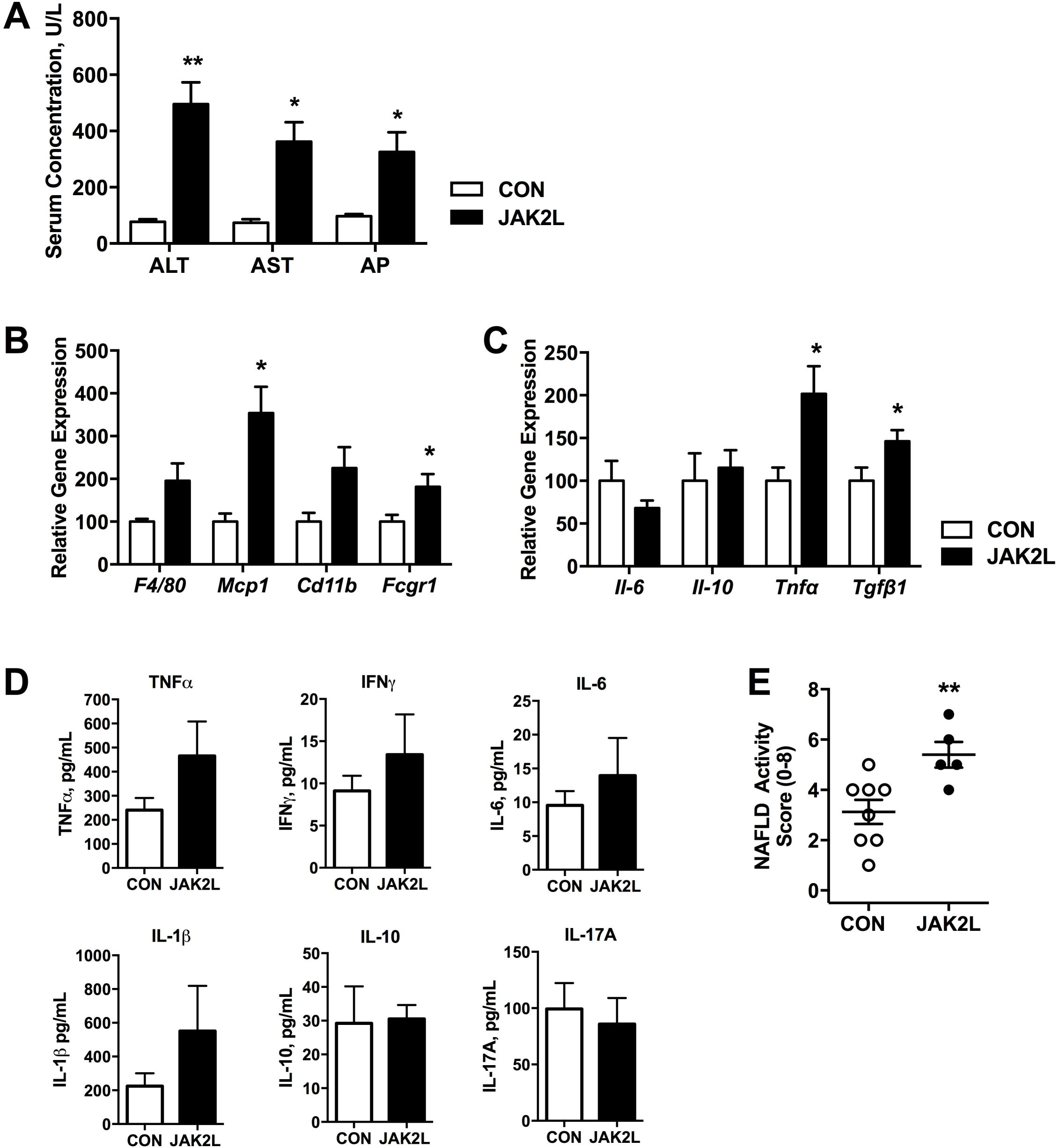
Markers of inflammation are elevated in JAK2L mice. (A) Serum markers of liver damage, Alanine transaminase (ALT), aspartate transaminase (AST) and alkaline phosphate levels are increased in JAK2L mice compared with Controls. (B) Relative gene expression of inflammatory cell markers and (C) inflammatory cytokines in livers of aged mice. (D) Serum levels of inflammatory cytokines Tnfα, interferon-γ (INF-γ), IL-6 and IL-1β, IL-10 and IL-17A. (E) Cumulative NALFD activity score (NAS) was calculated using scores for steatosis (0-3), lobular inflammation (0-2), hepatocellular ballooning (0-2). Data represent mean ± SEM, n=8 for Control and n=5 for JAK2L. * P < 0.05, ** P < 0.01

In NAFLD, recruitment of immune cells to the liver and up-regulation of pro-inflammatory cytokines incites an inflammatory state in the liver (39). Hepatic gene expression analysis of macrophage markers revealed higher expression of *F4/80*, *Mcp1*, *Cd11b* and *Fcgr1* in JAK2L livers as compared to controls (Figure 2B). Hepatic expression of inflammatory cytokines, interleukin (*Il)-6* and *Il–10* did not vary between the groups but *tumor necrosis factor-*α (*Tnf*α) expression was 2-fold higher in JAK2L livers (Figure 2C). However, circulating levels of Tnfα and other inflammatory cytokines including interferon-γ (INF-γ), IL-6, IL-10, IL-17A, and IL-1β were not significantly elevated in JAK2L mice (Figure 2D). Despite this, there was histological evidence of inflammation in these aged mice. The presence of lobular inflammation, micro-granulomas and portal inflammation was higher in JAK2L mice (Table 1). Also, markers of liver injury (acidophil bodies and pigmented macrophages) were more prevalent in JAK2L mice. Therefore, using these data we generated a NAFLD Activity Score (NAS) for each group, which takes into account scores for steatosis, lobular inflammation and ballooning and positively correlates with NASH (40). The average NAS for control mice was 3.13 as compared to 5.4 for JAK2L mice (Figure 2E) suggesting a higher propensity towards developing NASH in JAK2L mice.

### Aged JAK2L livers are markedly fibrotic

NASH progresses to liver fibrosis and overt cirrhosis in 10-15% of patients. Other models of impaired hepatic GH signaling have reported increased fibrotic gene expression (24). In our aged JAK2L cohort, liver collagen deposition was increased (Figure 3A). Hepatic fibrosis in JAK2L mice was a mix of periportal, centrizonal-chickenwire and septal fibrosis with an average fibrosis score of 2.3 for JAK2L as compared with 0.25 for control mice (Figure 3B). Other scoring methods to assess NASH including Brunt Staging and overall NASH scores were also significantly higher in the JAK2L cohort.

**Figure 3.**
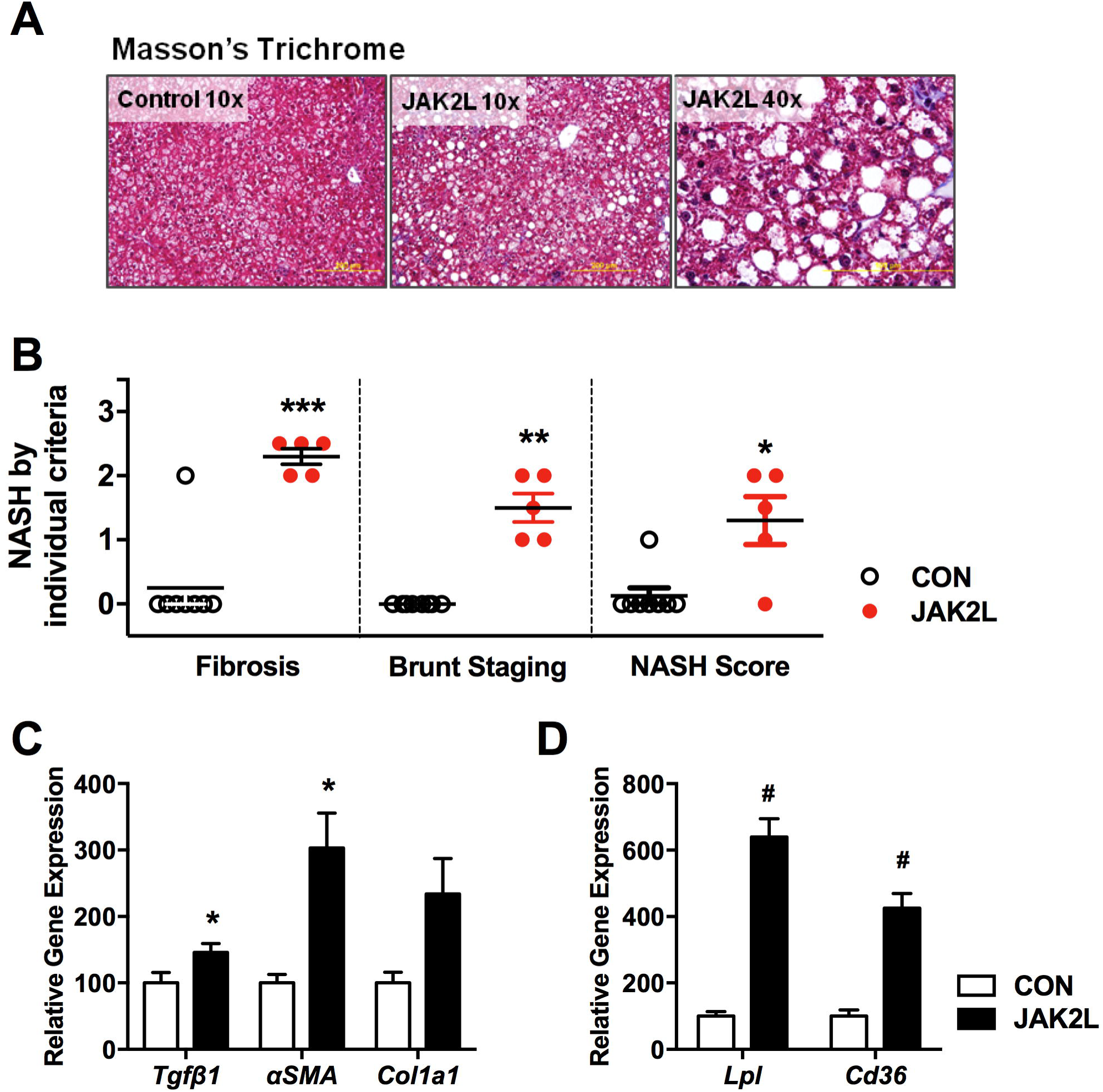
Older JAK2L mice develop NASH with fibrosis. (A) Masson’s trichrome staining of liver sections shows increased collagen deposition in JAK2L mice compared to Controls. (B) NASH scores by individual criteria: Fibrosis (0, none; 1, periportal; 2, centrizonal-chickenwire; 2.5, septal; 3, early bridging; 4, cirrhosis), Burnt staging (1-4) and NASH (0-4). Relative hepatic gene expression of (C) fibrosis markers and (D) NASH markers lipoprotein lipase and Cd36. Data shown as mean ± SEM, n=8 for Control and n=5 for JAK2L. * P < 0.05, ** P < 0.01, *** P< 0.001, # P< 0.001

Histological identification of fibrosis was supported by gene expression analyses. Higher expression of *Tgfβ1*, α-smooth muscle actin (*αSMA*) and *Col1a1* was present in JAK2L livers (Figure 3C). Additionally, lipoprotein lipase (*Lpl*) and fatty acid translocase (*Cd36*) were significantly increased over 6- and 4-fold, respectively, in JAK2L compared to control livers (Figure 3D). This is of particular interest as *Lpl* and *Cd36* are elevated in the setting of steatosis and NASH (41). Together these data demonstrate that the JAK2L mouse is model of spontaneous fatty liver that progresses to NASH with fibrosis.

### Mice with Jak2-deficient adipocytes are protected from developing age-associated NASH

We previously demonstrated that hepatocyte-specific loss of *Jak2* causes hepatic steatosis. Here, we showed that steatosis progressed to NASH in older JAK2L mice in a mixed background. Conversely, we have also shown that *Jak2* deletion in adipose tissue (JAK2A mice) protects against the development of fatty liver (36). In addition, mice with concomitant loss of *Jak2* in hepatocytes and adipocytes (JAK2LA mice) have ~60% less liver TG and a reduction in GH-stimulated lipolysis compared to their JAK2L littermates. Therefore, we next asked whether the development of NASH in JAK2L mice could be prevented in mice with concomitant adipocyte *Jak2* deletion (JAK2LA mice). To test this hypothesis, we crossed JAK2L mice to JAK2A mice expressing Cre recombinase under control of the adiponectin promoter as previously described to generate control, JAK2L (hepatocyte *Jak2* deletion), JAK2A (*adipocyte* Jak2 deletion) and JAK2LA (hepatocyte AND adipocyte *Jak2* deletion) mice on C57Bl/6 background (42). After 65-70 weeks of age, mice were analyzed for histological, genetic and biochemical markers of NASH.

H&E stained liver sections of the aged mice showed lipid accumulation in control, JAK2L and JAK2LA cohorts but almost no lipid droplets in JAK2A hepatocytes (Figure 4A). Trichrome stained sections also showed collagen deposition in JAK2L mice with minimal to no collagen in control or JAK2A mice and less in JAK2LA mice. Control livers had on average 47% of the hepatocytes with lipid droplets as compared to 80% in JAK2L mice, 1% in JAK2A mice and 14% in JAK2LA mice (Figure 4B). For control and JAK2L livers, steatosis was primarily localized to zone 3, whereas in JAK2LA livers, lipid-laden hepatocytes were also found in the azonal and panacinar regions. We also found an uneven distribution in the type of lipid droplets among the groups. JAK2L livers had significantly more microvesicular than JAK2A or JAK2LA (Figure 4C). Conversely, there was no statistically significant difference between the 4 genotypes in the prevalence of macrovesicular steatosis. TG and cholesterol levels were similar between the control and JAK2L cohorts (Figure 4D). Liver TG was significantly lower in JAK2A versus control and JAK2L mice, while cholesterol was reduced in both JAK2A and JAK2LA versus the control and JAK2L cohorts.

**Figure 4.**
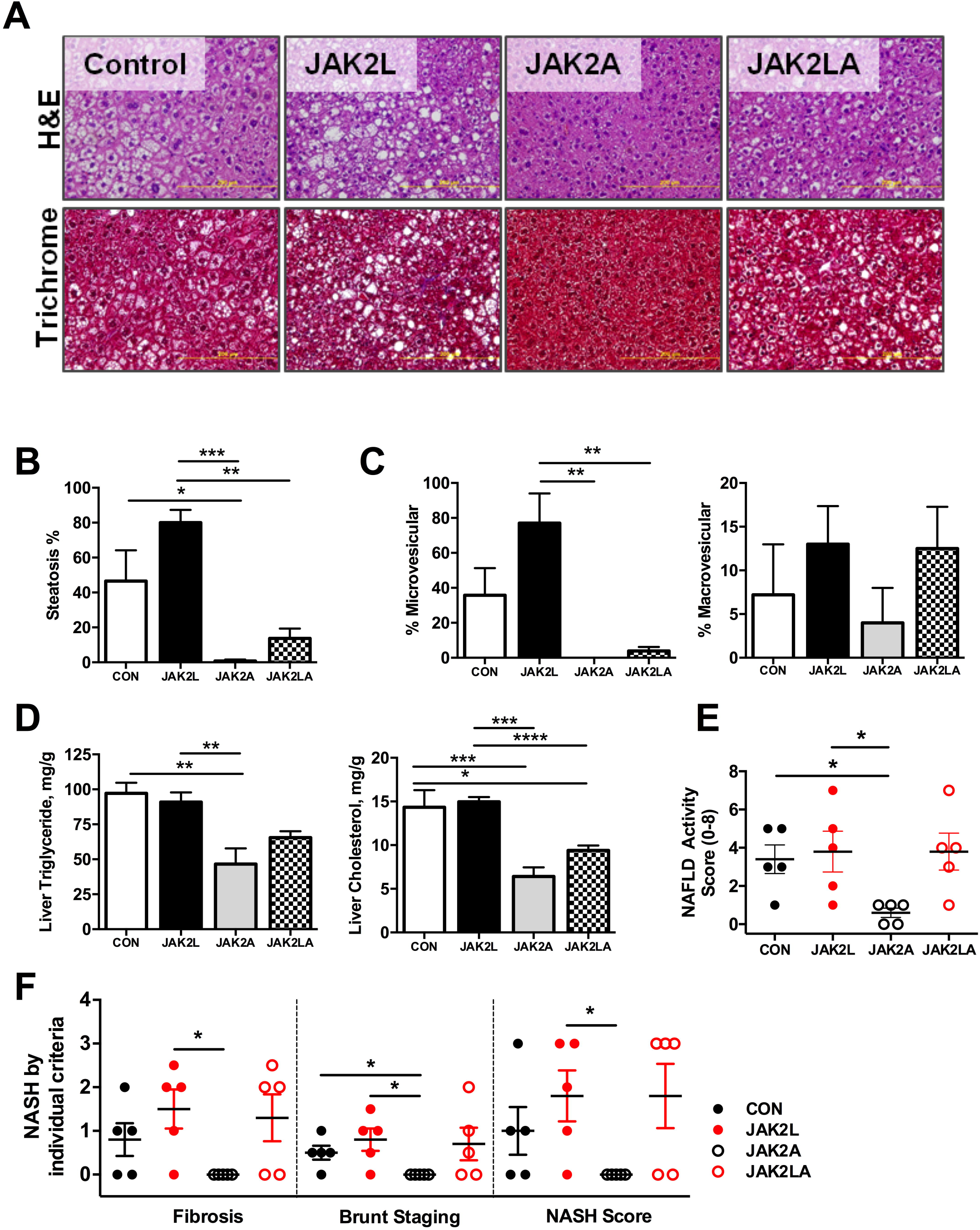
Jak2 deletion in adipocytes protects mice from developing NASH but not on the JAK2L background. Control, JAK2L, JAK2A and JAK2LA mice on C57BL/6 background were aged for 65-70 weeks. (A) Hematoxylin and eosin (H&E) (top) and Masson’s trichrome stain (bottom) of liver sections. (B) Steatosis percent and (C) steatosis type (micro- or macro-vesicular) was determined from H&E stained sections. (D) Liver triglycerides and cholesterol esters levels were reduced in JAK2A and JAK2LA mice. (E) NALFD Activity Score and (F) NASH scores. Data shown as mean ± SEM. N=5 per group. * P < 0.05, ** P < 0.01, *** P < 0.001, **** P< 0.0001.

We also measured the presence of inflammatory foci and liver injury (Supplemental Table 2) and found that there were similar levels of lobular and portal inflammation in control and JAK2L livers. There was a higher percentage of injured hepatocytes in JAK2L livers. JAK2A livers were the least inflamed and lacked hepatocyte damage. Surprisingly, a large percent of JAK2LA livers had inflammatory foci, portal inflammation and balloon cells. The average NAS scores were 3.4, 3.8, 0.6, and 3.8 for the control, JAK2L, JAK2A, JAK2LA cohorts, respectively (Figure 4E), demonstrating significant protection from steatosis and inflammation in JAK2A mice with almost no risk for development of NASH.

Histology scores for fibrosis, Brunt staging and overall NASH score were highest in the JAK2L and JAK2LA cohorts as compared to control animals and lowest in JAK2A mice (Figure 4F). Therefore, while loss of adipocyte *Jak2* on the JAK2L background reduced benign steatosis, it did not protect mice from developing NASH. These data suggest there is a liver autonomous component to the development of NASH in *Jak2*-deficient livers.

### Adipose Jak2 deletion does not protect JAK2L mice from hepatic inflammation

There was a trend toward increases in ALT, AST, and alkaline phosphate in JAK2L mice as compared to controls while levels were significantly lower in JAK2A versus JAK2L mice (Figure 5A). There were no significant changes in levels of inflammatory cytokines between the groups (Supplemental Figure 1). We measured genetic markers of steatosis and NASH. *Lpl* mRNA expression was 4-6 fold elevated in JAK2L and JAK2LA livers as compared to controls (Figure 5B). Consistent with our previous findings, *Ppar*γ and *Cd36* gene expression were also elevated in JAK2L and JAK2LA mice (21, 28).

**Figure 5.**
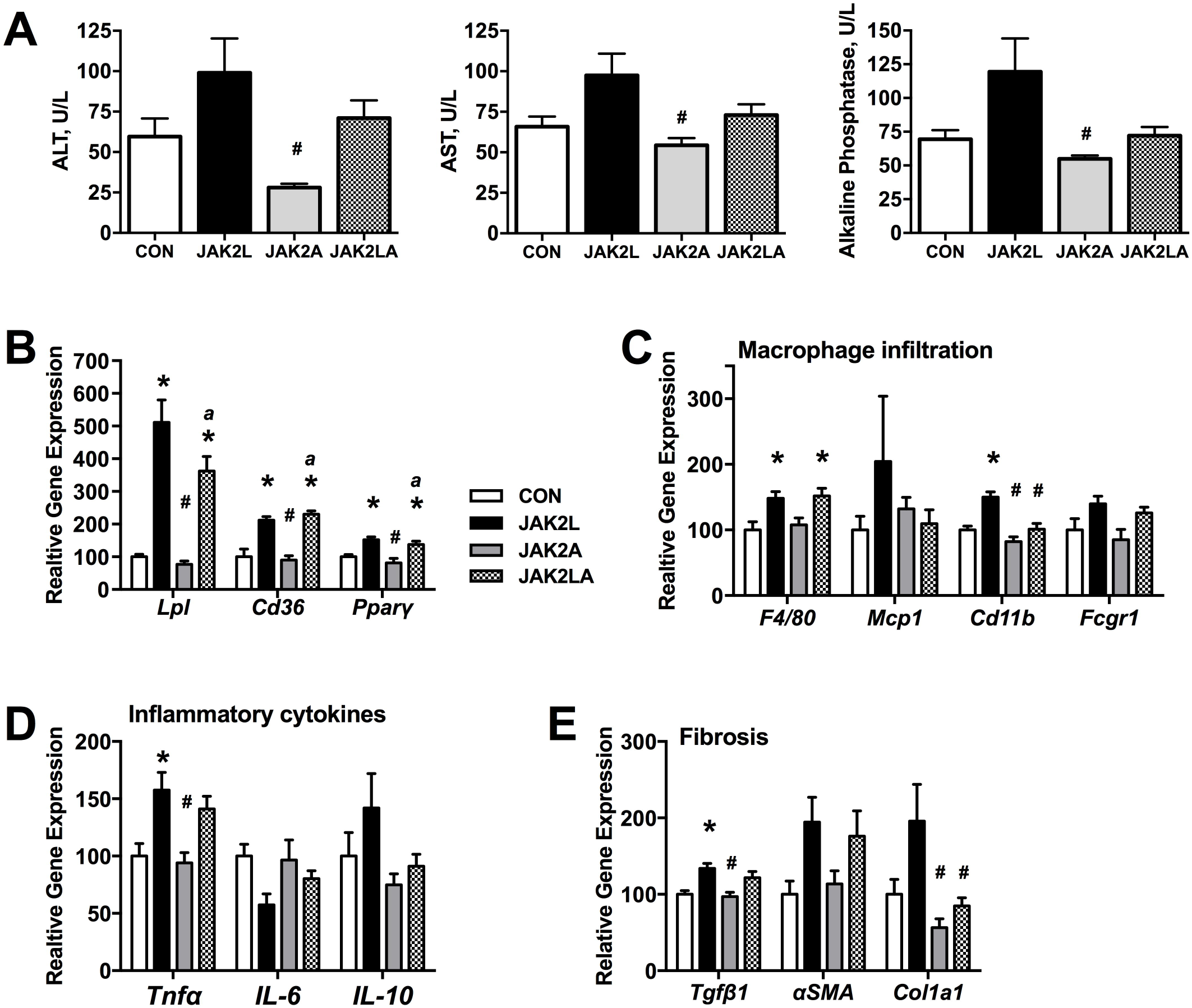
Hepatic inflammation is increased in JAK2L mice. A) ALT, AST and alkaline phosphate levels are increased in JAK2L mice. (B) Genetic markers of steatosis and NASH are elevated in JAK2L and JAk2LA livers. (C) Macrophage infiltration, (D) inflammatory cytokine and (E) fibrosis gene expression was elevated in JAK2LA mice. Data shown as mean ± SEM. N=5 per group. * P < 0.01 compared to CON, # P < 0.01 compared to JAK2L and a P < 0.01 compared to JAK2A.

Markers of macrophage infiltration were all elevated in JAK2L livers compared to controls while only *F4/80* and *Fcgr1* were elevated in the JAK2LA cohort (Figure 5C). As seen in the mixed background JAK2L mice, *Tnf*α expression was elevated in JAK2L and JAK2LA livers (Figure 5D). JAK2A mice had normal levels of all transcripts. Fibrosis-associated genes were increased in JAK2L livers compared to controls (Figure 5E) and α*SM-actin* was also increased in the JAK2LA cohort. Therefore, despite less steatosis, JAK2LA mice develop as much inflammation and hepatocyte damage as JAK2L animals and were not protected from developing NASH due to los of *Jak2* in adipocytes. This suggests that loss of hepatic Jak2 and/or GH signaling directly impacts the risk for developing NASH independent of lipolysis and demonstrates a dissociation between liver lipid and risk of NASH and fibrosis.

### Dissociation between insulin resistance, obesity, and NASH in mice with tissue specific deletion of Jak2

To understand potential extra-hepatic factors contributing to the development of NASH in JAK2LA mice we next examined body composition and insulin resistance. Previously we reported that adiposity is increased in JAK2A and JAK2LA mice due to reduced lipolysis (36). In this aged cohort, we also found that JAK2L mice weighed significantly less than controls while JAK2A and JAK2LA mice were the heaviest of the groups (Supplemental Table 3). As shown in Figure 6A, JAK2L mice had 6.6g of fat compared to 20.5g, 24.0g, and 19.0g for the control, JAK2A, and JAK2LA cohorts, respectively. As a result, the total fat mass contributing to body composition was 21%, 39%, 45%, and 45% for JAK2L, control, JAK2A, and JAK2LA mice, respectively (Figure 6A right). As such, the difference in body weight in these cohorts was primarily due to differences in body fat content, consistent with the known fat wasting effect of high GH levels in JAK2L mice and the reduced adipocyte lipolysis in JAK2A and JAK2LA animals. Furthermore, complete protection from steatosis in JAK2A mice demonstrates that adiposity per se does not increase the risk of metabolic liver disease.

**Figure 6.**
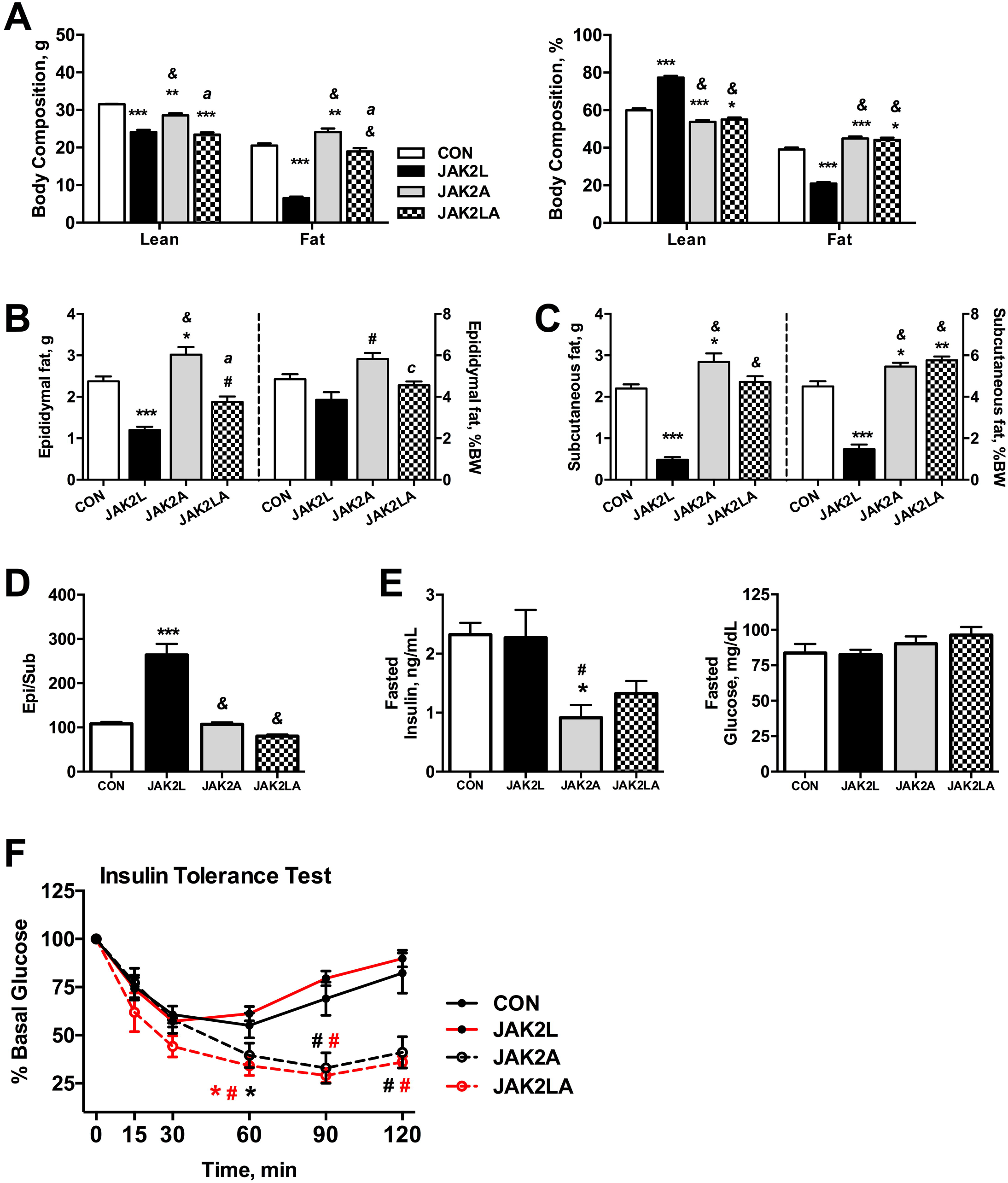
Insulin sensitivity is improved in JAK2LA mice despite obesity and NASH. (A) Body composition showing lean mass and fat mass expressed in grams or as a percent of total body weight. (B) Epididymal fat depot and (C) subcutaneous fat depots were smallest in JAK2L mice. (D) Ratio of epididymal fat weight to subcutaneous fat weight expressed ad a percent. (E) Fasted insulin and glucose levels in aged mice. (C) Insulin tolerance test was performed on mice by after a 4 hr fast. Data shown as mean ± SEM, N= 6-10 per group. * P< 0.05, ** P < 0.01 and ** P < 0.001 compared to CON. # P < 0.01 and & P < 0.001 compared to JAK2L, c P < 0.05, b P < 0.01 and a P < 0.001 compared to JAK2A

Examination of the individual fat depots showed that both epididymal and subcutaneous fat pad weights were positively correlated with fat mass (Figure 6B). However, the effect on subcutaneous fat was most striking with JAK2L mice having approximately 4-fold less than that of the other cohorts (Figure 6C). The epididymal to subcutaneous fat ratio was 1 in both control and JAK2A mice but almost 3 in JAK2L animals and less than 1 in the JAK2LA cohort (Figure 6D).

To determine the relationship between metabolic liver disease and insulin resistance, we next examined insulin sensitivity. Fasted insulin levels were similar in the control and JAK2L, but significantly lower in JAK2A and JAK2LA mice (Figure 6E). There was no difference in fasted glucose levels between the groups. To test whole body insulin sensitivity, we performed insulin tolerance tests. Control and JAK2L mice were the least insulin responsive of the cohorts (Figure 6F). JAK2A mice had a maximal 67% drop in blood glucose levels following insulin injection and surprisingly, JAK2LA mice were also very insulin sensitive. This suggests that in JAK2A and JAK2LA animals, loss of Jak2 in adipocytes augments whole-body insulin sensitivity. Notably, JAK2A and JAK2LA mice retained exquisite insulin sensitivity despite a 45% body fat composition. Conversely, despite leanness, JAK2L mice were relatively insulin resistant, demonstrating a strong dissociation between adiposity and insulin sensitivity. Further, in JAK2LA mice, insulin sensitivity was augmented despite the presence of hepatic steatosis, inflammation, and fibrosis. Collectively then, loss of hepatocyte *Jak2* activity may be the primary contributor to the development of NASH while adipocyte *Jak2* appears to mediate insulin sensitivity. Overall the JAK2LA mice represent a fascinating disassociation of metabolic liver disease and body composition from insulin sensitivity.

## DISCUSSION

Here, we describe a mouse genetic model with aging-related spontaneous progression from simple hepatic steatosis to NASH with fibrosis, insulin resistance, and low body fat. The simple steatosis previously described in JAK2L mice progressed to NASH with fibrosis and many key features of human NASH, including steatosis, hepatocyte damage and inflammation (3). Mice with disruption of *Jak2* from liver and fat (JAK2LA) mice developed less steatosis despite increased adiposity and insulin sensitivity, but they progressed to NASH, just as in JAK2L mice. This suggests that there may be a liver autonomous factor that promotes hepatic inflammation and increases risk for NASH in *Jak2*-deficient hepatocytes that is independent of lipolysis, adipocyte JAK2 signaling, and insulin sensitivity. Conversely, loss of adipocyte *Jak2* maintains and augments insulin sensitivity even in the face of obesity and NASH.

Our findings are not consistent with Shi et al (43) who reported no progression to NASH in their model of hepatocyte-specific Jak2 deletion. The discordance between our findings and theirs may be due to differences in age or the background strain of mice used. For example, we found that NASH was more severe in JAK2L mice on a C57BL/6,129/Sv mixed background as compared to a pure C57Bl6. Consistent with our findings, several others have reported a link between growth hormone deficiency and prevalence of NAFLD and NASH in both rodents and patients (22, 24, 44, 45).

We were surprised that there was no protection from NASH in the JAK2LA as compared to the JAK2L mice. JAK2L and JAK2LA mice are genetically identical in the liver. They both have near-complete reductions in circulating IGF1 and a compensatory increase in circulating GH. As we showed previously, JAK2LA mice have reduced GH-stimulated lipolysis and are largely protected from hepatic steatosis when on normal chow and at a young age (42). However, we have recently found that JAK2L and JAK2LA mice have similar levels of liver lipid when fed high fat diet (46). Since hepatic CD36 is elevated similarly in JAK2L and JAK2LA mice, we presume that the increase in dietary lipid on HFD leads to significant increased uptake of FFA and equalization of hepatic triglyceride content. This is consistent with what we observed here in the aged cohort on normal chow, where there is similar liver lipid content in JAK2L and JAK2LA mice. That is, adipocyte *Jak2* disruption appears to protect against hepatic steatosis when dietary lipid levels are low and at a young age. With increased dietary lipid or advanced age, there is no difference in liver lipid level between JAK2L and JAK2LA mice.

There is a striking difference in the relative degree of macro-versus microvesicular steatosis in the 70-week cohort, with a dramatic increase in microvesicular steatosis in JAK2L mice not seen in JAK2LA mice. The classification of rodent and human NASH is similar though there are not many reports of mice with predominantly microvesicular steatosis (34, 47, 48). In humans, microvesicular steatosis has been associated with defects in β-oxidation (48). We also observed a strong association between the degree of microvesicular steatosis and the degree of insulin resistance. That is the total amount of liver triglyceride, degree of macrovesicular steatosis, and measures of NASH and fibrosis are equal in JAK2L and JAK2LA mice. The only difference is between the degree of microvesicular steatosis and insulin resistance. We hope to explore this fascinating association in future work.

There are interesting differences and similarities between the JAK2A and JAK2LA mice. In the JAK2A cohort, there was complete protection from NASH <u>and</u> protection from aging-associated insulin resistance. In the JAK2LA cohort, there was protection from aging-associated insulin resistance <u>despite</u> progression to NASH. This suggests that insulin sensitivity does not afford protection from progression to NASH. This also suggests that in this model, insulin sensitivity does not protect from NASH nor does insulin resistance necessarily cause NASH and that NASH might not directly cause insulin resistance. Recently, we found that JAK2A mice are similarly protected from fatty liver and insulin resistance when challenged with high fat diet (46). In the present study, insulin sensitivity was maintained despite 45% body fat composition, demonstrating dissociation between obesity and insulin resistance.

Most strikingly, JAK2LA mice also maintained insulin sensitivity despite increased adiposity and significant NASH and liver fibrosis. Increased insulin sensitivity due to adipocyte Jak2 deletion appears to be adipocyte autonomous. One of the key differences between the cohorts is the lipolytic activity of adipose tissue. JAK2A and JAK2LA mice have decreased lipolysis and JAK2L mice have increased lipolytic activity (20). Others have shown that inhibiting lipolysis in healthy subjects reduces FFA flux and improves liver function and insulin sensitivity without affecting hepatic fat content (49). Therefore, it is possible that lipolysis and increased FFA flux is the driver of whole body insulin sensitivity but not NASH progression.

In summary, here we present data on a genetic model of spontaneous NAFLD with progression to NASH and fibrosis. The liver pathology is at least partially independent from systemic metabolic changes in body composition or insulin sensitivity. To our knowledge, this is the first such model to be reported and represents a significant finding that disrupts the dogma that metabolic liver disease and systemic metabolic disease are inextricably linked.

